# Quantifying optical sectioning in reflection microscopy with patterned illumination

**DOI:** 10.64898/2026.02.06.703262

**Authors:** Cathie Ventalon, Agathe Nidriche, Delphine Débarre

## Abstract

Sectioning techniques based on patterned illumination have been widely used to obtain well-contrasted images of thick samples using widefield imaging setups. While their application to fluorescence microscopy has been extensively demonstrated and studied, their application to reflection imaging is scarcer and their performance has only been partly characterized. In this paper, we study numerically and analytically two such sectioning techniques, line confocal (LC) and structured illumination microscopy (SIM), in the context of their application to coherent reflection imaging. We derive approximate analytical equations to relate the performance of sectioning to the optical setup parameters, allowing straightforward understanding of their influence on the image intensity and depth of focus, and we systematically compare our predictions with experimental data. Finally, we quantify the precision and accuracy of each method in typical practical cases, providing guidelines to choose the most appropriate (LC, SIM, or a simple background subtraction on a widefield image) for the sample under study. We illustrate optical sectioning in the particular case of reflection interference contrast (RIC) microscopy, an imaging technique widely used in soft matter and biophysics studies to monitor object-surface interactions, or quantify surface functionalization. Our derivation, however, should also prove useful for other reflection methods such as optical coherence tomography (OCT) or flood illumination ophtalmoscopy.

## 1. Introduction

Imaging methods based on reflected light have proven valuable in a variety of biological contexts due to their ability to provide structural images based on endogenous contrasts. A number of these techniques use a widefield implementation that permits fast frame rates and minimize motion artifacts. When imaging thick samples, however, their sensitivity is hampered by incoherent reflections from out-of-focus structures that limit their contrast and the possibility to quantitatively analyse image intensity information. In this context, optical sectioning techniques based on the modulation of the illumination pattern have been established that mostly maintain fast imaging frame rates: structured illumination microscopy (SIM) [1, 2] and line confocal (LC) [3] microscopy. In the present article, we aim to derive numerical and analytical predictions for the performance of each of these approaches that provide a direct link with the parameters of the optical system.

Indeed, despite having been considered in a number of articles over the past 30 years for fluorescence imaging [4–8] or to a lesser extend for reflected light [1, 9], the problem of axial sectioning obtained by optical patterning has received mostly complex treatments incorporating high-numerical aperture effects. In addition, depth of focus is usually ill-defined in (partially) coherent systems as it depends on the structure of the sample, which complexifies the derivation of simple, approximate formulas for the axial resolution and detected intensity. While the published approaches provide exact or very accurate predictions for the axial sectioning ability, they are not always easily translated into guidelines for adjusting setup parameters or comparing different sectioning techniques. To complement these existing theories, we derive here analytical expressions for the axial sectioning obtained using either LC or SIM with (partially) coherent imaging and relatively low illumination numerical apertures (INA<1).

This particular framework has been chosen to mimic representative experimental conditions for a particular reflection contrast imaging method: reflection interference contrast (RIC) microscopy, also known as interference reflection microscopy (IRM). This technique records the interference pattern created by the light reflected (or scattered for nanoobjects) by two or more well-defined or diffuse optical interfaces located within the coherence volume near the focus of a wide-field microscope. This interference pattern encodes information about the optical distances between the interfaces, which can be quantitatively analyzed with appropriate modelling [10, 11]. It has emerged as a powerful method for the characterisation of thin organic hydrated films [12, 13], interface phenomena such as wetting, contamination or crystal growth [14], as well as quantitative studies of cell adhesion [15, 16]. We have recently demonstrated the use of SIM and LC to specifically detect the RIC signal in the vicinity of the focal plane, thereby increasing the contrast of interferences and allowing quantitative analysis of the reflected intensity [17]. However, the derivations presented in this paper apply to any coherent reflection techniques such as optical coherence tomography (OCT) or flood illumination ophtalmoscopy (FIO).

Here, we first introduce our experimental setup and the modelling assumptions used in the subsequent derivations (section 3.1). We then derive approximate theoretical expressions for the image intensity and optical sectioning, and compare them with more accurate numerical predictions (section 3.2). We confirm our theoretical predictions by comparison with experimental results on a model sample (section 3.3). Finally, we discuss the noise and remaining spurious signal in the sectioned images for several typical samples, which permits comparing the relative benefits and limitations of LC and SIM sectioning in the context of coherent reflection imaging, and we illustrate these benefits on example cases of RIC imaging of thin films and cells (section 3.4).

## 2. Methods

### 2.1. Optical setup

The optical setup has been fully described elsewhere [17]. Briefly, it is based on a modified IX71 Olympus inverted widefield microscope, with home-built system separating illuminating from reflected light based on a polarisation beamsplitter cube (PBS251, Thorlabs) and a custom-made optically flat achromatic quarter waveplate (Fichou, France). Illumination is provided by an incoherent plasma light source (HPLS345, Thorlabs) that is spectrally and spatially filtered using a dichroic filter (FF01-457/530/628-25, Semrock) and graduated diaphragms (SM1D12C, Thorlabs). Before filtering, illumination light is reflected off a high-speed digital micromirror device (DMD - V-9601, Vialux, Germany) conjugated with the image plane of the microscope objective (depending on the experiment and as mentioned in the text: UPLSAPO60XO, UPLSAPO20XO, and UPLSAPO20X Olympus, Japan). Detection is achieved on a sCMOS cameras (ORCA-Flash4.0 V3, Hamamatsu, Japan) after filtering with a dichroic filter (FF01-452/45-25, FF01-530/43-25 or FF01-629/56-25 depending on the detected wavelength, Semrock). For LC imaging the DMD and the sCMOS are synchronized using a pulse generator (4052 waveform generator, BK precision). The magnification between the DMD and camera planes is chosen such that one DMD pixel (10.8 µm) is projected onto two camera pixels (2 × 6.5 = 13 µm, corresponding to M_*l*_ = 1.2). Details about the hardware synchronization, image acquisition program and optical alignment are given in [17].

### 2.2 Sample preparation

Gold coating was performed on a home-built sputtering deposition machine. Glass coverslips (n°1 type, thickness ∼150 *µ*m, Thermo Scientific Menzel) of 32 mm diameter were piranha-cleaned and used as substrates. A ≈0.5 nm titanium layer was first deposited to prevent gold dewetting, before ≈5 nm gold was added.

Poly-N-isopropylacrylamide (PNIPAM) brushes were grown and patterned as described elsewhere [18]. The brush dry thickness was characterized using a home-built ellipsometer and an identical brush grown on a piece of silicon wafer.

Red blood cells were obtained from a healthy donor through the French Blood Bank (agreement EFS AURA 22-051) and prepared as described elsewhere [19]. The glass coverslips (30 mm diameter, VWR International, Radnor PA, USA, 631 − 0175) were treated with plasma for 15 s before deposition of 500 *µ*L 1 mg/mL Poly-L-Lysine (PLL) solution in ultrapure water (molecular weight 70 − 150 kDa, P1274-500MG lyophilised powder, Sigma-Aldrich). After 10 min incubation, the coated substrate is rinsed and dried under sterile conditions for a minimum of 2 hours. 2 mL of a freshly-rinsed-RBC suspension with a concentration of approximately 0.2 g/ L was then deposited on the coated coverslip shortly before measurements.

Image processing and analysis was performed using FIJI. All numerical calculations based on the equations derived below, as well as data processing and plotting, were performed using Mathematica.

## 3. Results

### 3.1. Coherent reflection imaging setup and modelling assumptions

The principle of our experimental setup is illustrated on Fig. 1: it is based on a widefield inverted microscope used in reflection mode, in which polarisation optics have been inserted in the turret to separate incoming and reflected light. Importantly, like in OCT or FIO and other so-called coherent techniques, the coherence of the light beam is not infinite but precisely controlled, both spatially (through an aperture diaphragm) and spectrally (through spectral filtering of the light source). This ensures that light is coherently reflected around the focal plane, while reflections from other planes do not produce unwanted interferences that would prevent analysis, and add incoherently to the total signal instead.

**Fig. 1.**
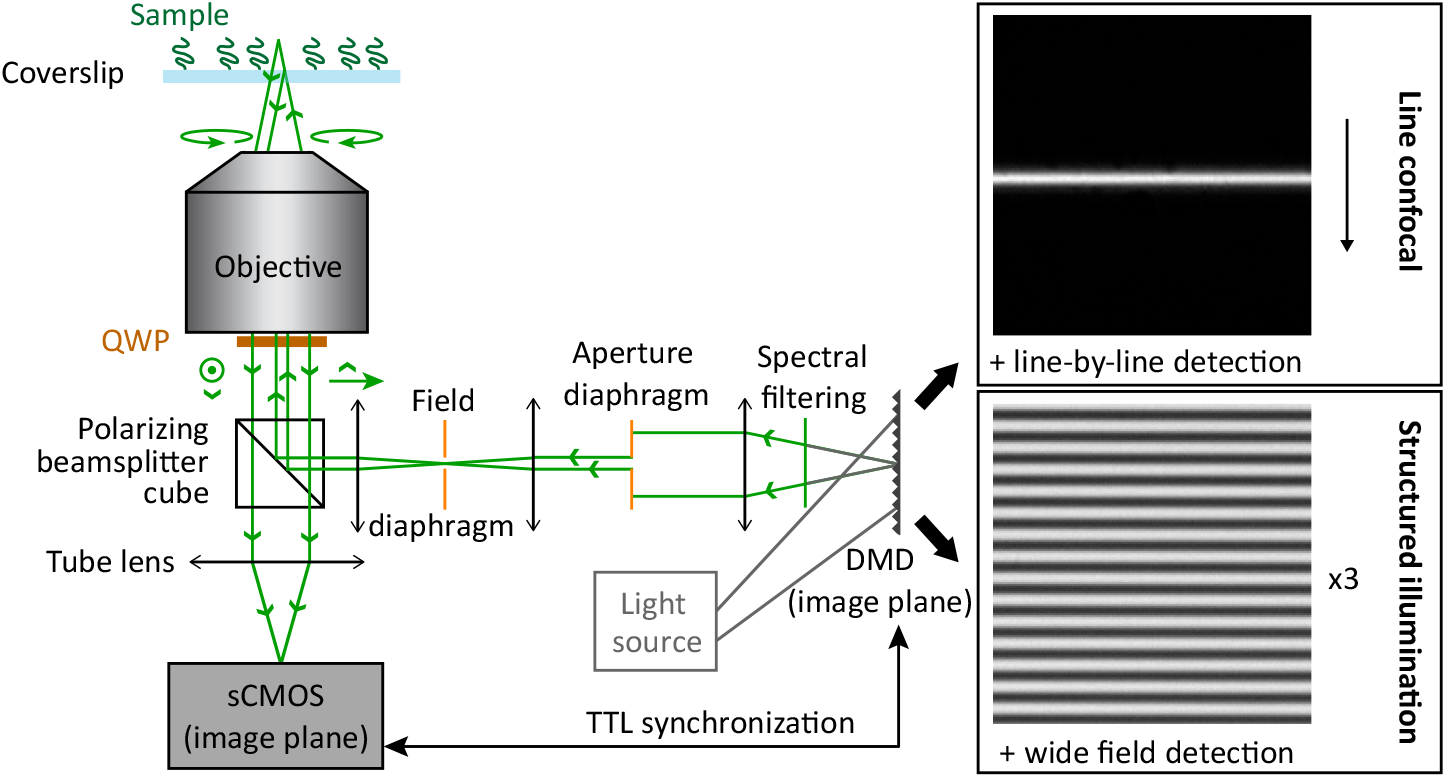
Schematic view of a RIC microscope with structured illumination. An incoherent white light source illuminates a digital micromirror device (DMD) with a numerical aperture large enough to neglect the subsequent diffraction by the DMD and related chromaticism. After spatial and spectral filtering setting the coherence of the illumination beam, the DMD is imaged in the image plane of the objective, which is itself conjugated with the sCMOS camera. Incoming and reflected light are separated using polarisation optics (a polarizing beamsplitter cube and a quarter waveplate, QWP; the polarisation of the beam is indicated with arrows at various positions in the light path). For simplicity, only one detected wavelength is shown here, but RIC images at different colours can be separated and imaged on different cameras or different parts of a single sensor using a spectral beamsplitting system after the tube lens. Insets: patterns displayed by the DMD for line confocal imaging (top) and structured illumination imaging (bottom).

Spatial coherence defines the illumination focal volume, setting both the minimal spatial period of illumination patterns that can be imaged on the sample, and the depth over which the smallest of these features blur with the distance from the focal plane. It is therefore the relevant scale for optical sectioning, as will be demonstrated in the next sections.

In contrast, the coherence volume, in which interferences can take place, combines both spatial and temporal coherence [11]. It is therefore at most as thick as the focal volume. As a consequence, interferences between reflections within the coherence volume are always located within the optically sectioned volume, and are not affected by optical sectioning (scaling with the focal volume). Instead, optical sectioning affects the relative intensities of the signals stemming from the focal volume compared to that stemming from structures further away in the sample that create an incoherent background. In the following, we will therefore not discuss interferences specifically, as these only modulate the in-focus signal of interest in a similar way (from the optical sectioning point of view) as a change in the reflection coefficient of a single in-focus interface.

Finally, RIC imaging is usually performed at a limited illumination numerical aperture (as set by the aperture diaphragm), INA: it is kept smaller than the full numerical aperture, NA, of the objective lens, to optimise the contrast of interferences. This is the definition of partially coherent imaging, where the spatial coherence of the illumination path is larger than or at most equal to the spatial coherence of the detection path. In the following, we will thus refer to “coherent reflection imaging” when the condition INA < NA is satisfied.

In addition to a classical reflection microscope, the setup incorporates a digital micromirror device (DMD) optically conjugated with the focal plane that permits a binary modulation of the illumination beam at high frequency (up to 20 kHz). This modulation is synchronized with image acquisition on a sCMOS camera. For LC imaging, a line of chosen width is scanned across the field of view and the signal on the camera is acquired progressively across the sensor within a matching moving line of fixed width [20, 21]. In this case, the defocussed reflections of the illumination line are rejected by the acquisition pattern as they fall mostly out of the moving detection line, which provides optical sectioning. Alternatively, widefield illumination and detection are used in SIM, and optical sectioning is achieved by demodulation of the imprinted grid illumination pattern through a combination of three images with a pattern shifted by 1/3 of its period. Numerous other implementations of SIM have been demonstrated [1, 22–26], but we restricted ourselves to this canonical version that permits quasi-real-time display of the sectioned image [17].

To model the optical sectioning obtained in these two configurations, we first introduce a number of assumptions which are of particular relevance for RIC imaging. First, in RIC microscopy, one prominent structure providing signal is the coverslip interface in contact or close vicinity with the sample under study (e.g. a polymer or protein layer, a cell or lamellar vesicle boundary, etc.). It is therefore relevant to consider as a typical sample a homogeneous plane interface. Similarly, in many cases, significant spurious reflections arise from other flat interfaces in the sample (liquid/PDMS and PDMS/air interfaces in a microfluidic channel, liquid/air interface in case of a free surface) that are best described as plane interfaces. Relatively large objects such as cells also locally act a a reflective plane. While this is not the case of a diffusing medium, such situation is rarely encountered in RIC imaging and is not specifically considered here. In the following, we will always consider a homogeneously reflective interface located between two media of the same refractive index. While this of course does not fully describe the properties of images on more complex samples (e.g. object curvature effects, scattering) which cannot be established as generally as for incoherent techniques, it provides an excellent approximation of the sectioning ability of the considered imaging setup. Reflective layers encountered in the retina or the surface of a tissue would also be well described in such a framework.

In addition, assuming that imaging takes place at or near the glass coverslip supporting the sample, we will neglect aberrations and diffusion in the light path.

Finally, RIC imaging is usually performed at relatively small INA (as set by the aperture diaphragm) to facilitate the quantitative analysis of interferences. This is in contrast with most uses of optical sectioning techniques, for which the best possible resolution is sought and where high numerical apertures are therefore preferred. Typically, the illumination numerical aperture is often kept smaller than 0.5 [11]. In this context, we will neglect polarisation effects associated with large illumination angles, as well as the associated variations in reflection coefficients of interfaces, and we will use the paraxial approximation to simplify the modelling of the optical system. While higher numerical apertures are used in some advanced implementations or other imaging methods, as well as in our experimental characterisations, we will stick to the simplified framework permitted by a low INA and will justify this choice a posteriori by comparison with experimental results.

### 3.2. Numerical and analytical modelling

In the following, we will start by establishing general equations describing the image formation process for line confocal and structured illumination coherent reflection microscopy within the above assumptions. The intensity and axial depth of focus will be numerically computed for a number of parameters. As a second step, we derive simple analytical approximations that are compared with the previous numerical calculations and provide simpler-to-use equations for relating the setup parameters to the achieved microscope performances.

For both LC and SIM, sectioning relies on the projection onto the sample of a pattern *P*(*x, y*) based on one or a series of parallel lines of light. We will therefore focus at first on the projection of a single line, described as:

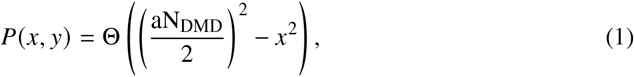

where Θ() is the unit step function, a is the size of a DMD pixel and N_DMD_ is the number of DMD pixels forming the width of the illumination line (Fig. 2(a)).

**Fig. 2.**
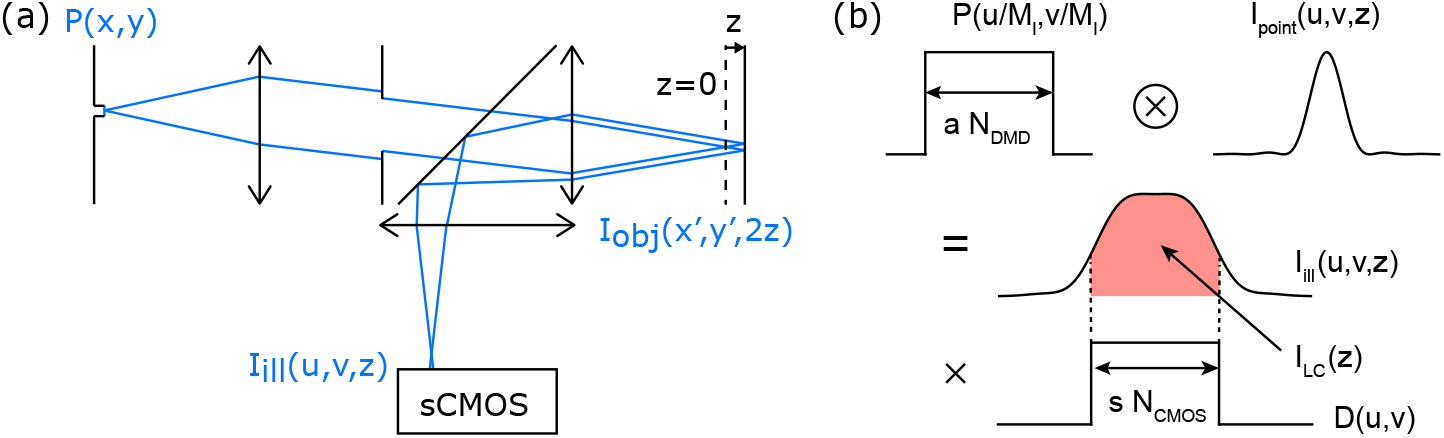
Scheme of line illumination for coherent reflection microscopy. (a), illumination scheme for a single projected line. The line pattern *P*(*x, y*) is projected onto the sample through an aperture diaphragm setting the illumination numerical aperture, INA. The intensity pattern at the focal plane, I_obj_(*x* ′, *y*′, *z*) depends on the distance between the focal plane and the reflective plane used as model sample, *z*. This intensity pattern is then imaged onto the camera through the full numerical aperture of the objective lens, producing the scaled pattern I_ill_(*u, v, z*). (b), detection scheme in LC coherent reflection microscopy. The illumination pattern on the camera, I_ill_(*u, v, z*), is obtained by convoluting the scaled line pattern, 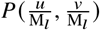, with the coherent transfer function I_point_(*u, v, z*). The detected signal, I_LC_(*z*), is then obtained by integrating this intensity profile over the detection function *D*(*u, v*).

This line is illuminated by an incoherent light source (both spatially and temporally), such that description of the image formation process is achieved by incoherent summation of the intensity pattern obtained for each of the points of this illumination line *P*(*x, y*). In contrast, for coherent reflection imaging the illumination pattern on the focal plane obtained from each of these points, I_obj_ (*x*′, *y*′, *z*), is obtained from the coherent transfer function of the optical setup, with *z* the displacement of the reflective plane with respect to the focal plane [27]. Using the paraxial approximation and neglecting high-angle polarization effects, we thus obtain:

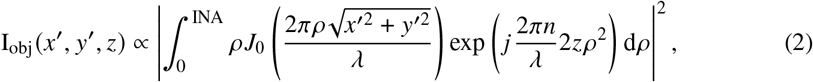

where *J*_0_ is the zeroth order Bessel function, λ is the wavelength of the illumination light, *n* is the refractive index of the medium filling the space between the reflective plane and the focal plane, INA is the illumination numerical aperture which has to be smaller than the full numerical aperture of the objective, and *j* ^2^ = −1. Note the multiplying factor 2 in the phase term that accounts for the reflection, such that a geometrical displacement 𝓏 of the reflecting plane results in an additional 2𝓏 path length for the reflected light. Because INA is kept smaller than NA, there is no additional aperture filtering on the detection path and the total instrument transfer function is equal to the coherent transfer function of the illumination path:

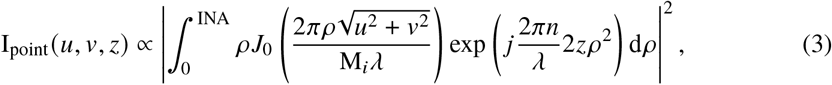

where M_*i*_ is the magnification factor between the image plane and the camera. From the two equations above, we find that the illuminated pattern on the camera for a single illumination line writes as:

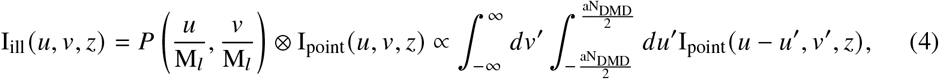

with ⊗the convolution operator and M_*l*_ the magnification factor between the DMD and the camera. Such expression can be used as a basis for the description of the two sectioning processes under consideration in this paper, through processes that differ in both cases. We will first consider the case of LC, while SIM will be addressed in the subsequent subsection.

#### LC coherent reflection image formation

In LC imaging, a single line is projected onto the sample and progressively scanned across the field of view. In a synchronised manner, a detection window in form of a line of thickness s ×N_CMOS_ (with s the camera pixel size and N_CMOS_ the detection line width in number of pixels) is swiped across the camera sensor to reconstruct an image through a 1D scanning process that, using a sCMOS camera, can be done at the same frame rate as full field imaging (see Fig. 1 and Methods). Matching the illumination and detection lines through careful synchronization and optical conjugation is key to efficient detection and provides optical sectioning through rejection of out-of-focus light falling outside of the detection line. A detailed description of the practical implementation of the method is provided in the supplementary data of [17].

In the following, we will assume that the detection line and the illumination line are perfectly synchronized, and that the acquisition frame rate is kept constant when changing the detection line width (i.e., the integration time per pixel is proportional to N_CMOS_). In this configuration, the temporal aspect of scanning and detection can be replaced by a static approach in which the detected intensity on each pixel is obtained by integrating over a spatial (instead of temporal) gate on the camera 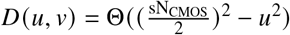. The detected intensity now only depends on the defocus 𝓏 and can be written as (Fig.2(b)):

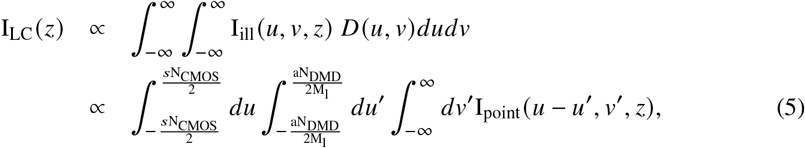

This equation allows full exploration of the impact of the different system parameters on metrics describing the quality of optical sectioning. To quantify the latter, we will consider in this study the image intensity at the focus, I_LC_ (0), and the full width at half maximum (FWHM) of the detected intensity as a function of defocus, *δ*_LC_, that we call depth of focus (Fig.3(a)). For the numerical calculations presented in this paper, we have set some parameters equal to their value in our experimental system: *a* = 10.8 *µ*m, *s* = 6.5 *µ*m, M_*l*_ = 1.2, *s*/ *a* = M_*l*_/2 = 0.6, M_*i*_ = 60, λ = 0.5 *µ*m and INA=0.5 with the refractive index of air. With these values fixed, we have studied the influence of N_CMOS_ and N_DMD_ on optical sectioning.

**Fig. 3.**
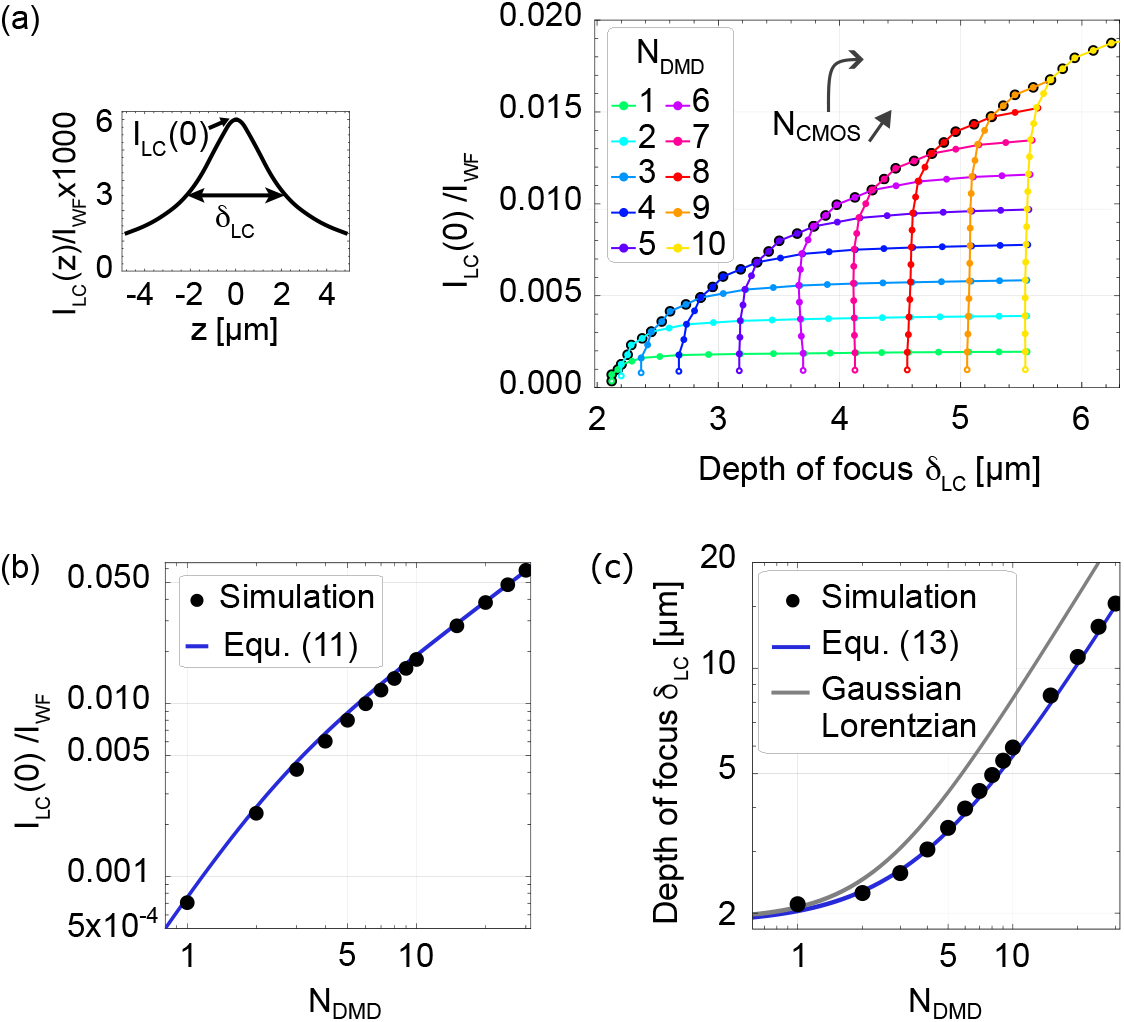
Line confocal sectioning in coherent reflection microscopy: numerical and analytical analysis. (a), left, example intensity profile in LC coherent reflection microscopy, defining graphically the parameters I_LC_ (0) and *δ*_LC_; right, simulated I_LC_ (0) /I_WF_ plotted against *δ*_LC_ as a function of the number of simultaneously detected lines N_CMOS_ (single colour, starting from empty marker) and of the number of illuminated lines N_DMD_ (colour-coded). The lines are a guide for the eye. Optimal (intensity, resolution) pairs are outlined in black and are obtained for N_CMOS_ ≈ 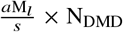. In (b-c), only this matching condition is plotted. (b), simulated normalised intensity as a function of N_DMD_ extracted from (a) (black dots), and theoretical predictions from Equ. 11 (blue line) showing excellent agreement. Both are normalised to the full field case, corresponding to N_tot_ = 1024 sCMOS lines in our setup. (c), simulated depth of focus *δ*_LC_ as a function of N_DMD_ extracted from (black dots), and theoretical predictions from Equ. 13 (blue line) showing also excellent agreement. Theoretical prediction using the classical Gaussian-Lorentzian approximation overestimates the depth of focus (gray line).

Fig.3(a) shows the image intensity vs. the LC depth of focus for various (N_DMD_, N_CMOS_) pairs. Each color line is obtained for a different line width on the DMD (see legend), and each dot of the curve corresponds to an increment of 1 on N_CMOS_, starting from N_CMOS_ = 1 at the bottom left (empty markers). Image intensity at the focus has been normalized to the widefield case (I_WF_) considering a similar illumination intensity in both cases. On such graph, the best performance is achieved for pairs corresponding to the highest intensity and smallest axial width, outlined in black on the graph. For each value of N_DMD_, these points are centered on 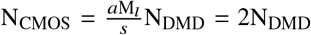, corresponding to matched widths for the illumination and detection lines in the camera plane, as is observed in confocal microscopy [8]. To optimize image quality, we will restrict ourselves in the following to this optimal setting so that only N_DMD_ will be considered a free parameter. Under these conditions, Fig.3(b) and (c) show respectively the intensity at focus, I_LC_ (0), and the depth of focus, *δ*_LC_, as a function of the line width (black dots): as expected, increasing N_DMD_ results in a brighter image, but at the expense of an increased depth of focus that reduces progressively the benefit of LC compared to full field imaging.

While the numerical assessment of LC optical sectioning properties is interesting to explore the relevant parameters for image quality, it does not provide a direct link between the system parameters and its performance (I_LC_(0) and *δ*_LC_, long range decay of the signal with defocus), which remains challenging to establish. We will therefore complement these calculations with a simplified approach to derive an analytical expression for I_LC_ (𝓏) (still normalised to its wide field counterpart) using relevant approximations. Again, we will consider only limited INA values commonly used in RIC microscopy. The beam shape obtained with a top-hat profile in the Fourier plane differs from the Gaussian-Lorentzian profile, and can instead be obtained from well-known integral expressions [9] Here, however, we are only concerned with the integrated profile along one spatial dimension, 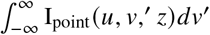, corresponding to the incoherent addition of point sources along the line direction (see Equ. 4 and 5). By fitting the exact beam profile integrated of *v*, we found that it is actually well approximated by a Gaussian-Lorentzian function, due to the smoothing of the side rings stemming from the top-hat profile edges:

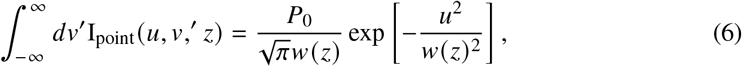

where *P*_0_ is the illumination power linear density, and *w*(𝓏) is the beam size at position 𝓏 from the focus. Here, however, the classical expression for *w*(𝓏) is empirically replaced by:

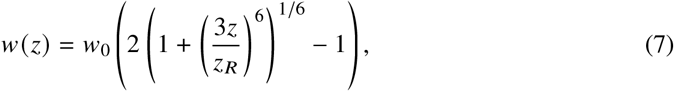

with 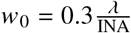 the intensity beam waist and

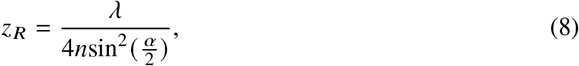

where *α* = arcsin (INA/ *n*) . Note that this description takes into account the factor 2 in 𝓏 stemming from reflection. Combining Equ. 5 and 6, we obtain by successive integration:

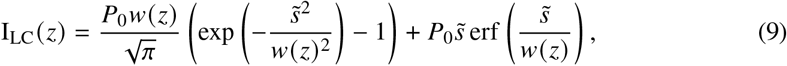

With 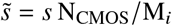. The above equation is very well fitted by the simpler expression:

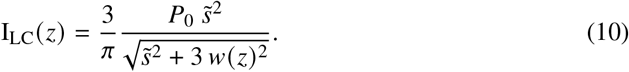

This can be understood as the product of the line width 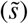, to which the collected signal is proportional, and a second term, 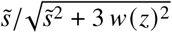, corresponding to a collection efficiency: this efficiency scales as the ratio of the detection line width by an effective width of the illumination pattern, itself given by the line width 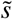 broadened by the beam size *w* (𝓏) at position 𝓏. Using this approximate expression one can extract simple analytical expressions for the image intensity at focus:

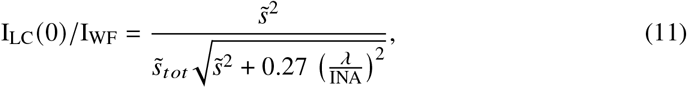

with 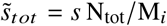 the total width imaged in the focal plane, and I_WF_ the intensity of the widefield (WF) image; and for the depth of focus:

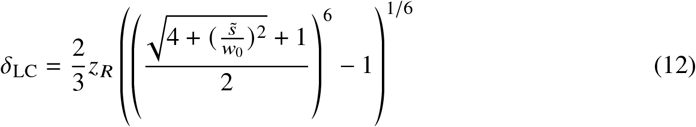

While this second expression is relatively complex, it can be approximated with a few percent error by the simpler expression:

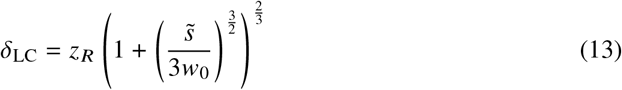

Equ. 11 and 13 match quantitatively the numerically-calculated values obtained from Equ. 5 (Fig.3(b) and (c), normalised to the brightfield intensity (with a total number of lines on the camera N_tot_ = 1024 in our setup)) while providing direct information on the optical sectioning properties of the system and their dependence on the system parameters. First, we retrieve with Equ. 10 the well-known slow ∝1/ (𝓏) signal decay away from the focus for I_LC_ 𝓏 that stems from confining illumination in one direction only, contrary to point-scanning confocal microscopy [4, 28]. Second, we observe that the depth of focus *δ*_LC_ follows a hyperbolic variation with the line width (a behaviour that has already been described in confocal microscopy [29]), while the image intensity grows mostly linearly with the same parameter: from this, we deduce that the optimal line width providing close-to-best optical sectioning while avoiding excessive reduction in image intensity is given by 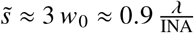. For this value, one obtains 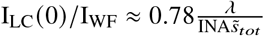 and *δ*_LC_ ≈1.59𝓏_*R*_. With our simulation parameters, this corresponds to I_LC_ (0)/ I_WF_≈ 0.006 and *δ*_LC_ ≈3 *µm*. Of note, using an accurate description of the beam size as a function of 𝓏 is essential to match the numerical simulations: the classical Gaussian-Lorentzian approximation, in which Equ. 7 is replaced by 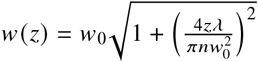, notably overestimates *δ* _LC_ .

#### SIM coherent reflection image formation

Many SIM schemes have been demonstrated since the first seminal implementation by Neil et al. [1]. Here, we have chosen to focus on the simplest and most common implementation which uses a 1D sinusoidal modulation of the illumination profile. Often (as in our experimental setup), however, this pattern is not directly imprinted on the illumination beam but a stripe pattern of alternating dark and bright lines is used instead due to the binary nature of DMD intensity modulation, and the sine modulation is in fact the result of apodization by the finite INA of this square modulation. The projection of this pattern onto the sample can thus be calculated numerically from Equ. 4 by incoherently adding the illumination pattern produced by a periodic assembly of several lines. Considering the classical SIM scheme, three images acquired with patterns shifted by 1/3 of their period (here referred to as I_1_, I_2_, I_3_) are then used to calculate a sectioned image through their pixel-per-pixel standard deviation:

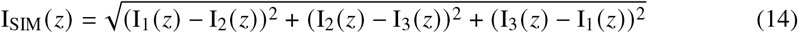

While more efficient algorithms may be used to minimise the image noise and improve its resolution, we have restricted ourselves here to the canonical variance calculation approach that has the benefit of simplicity and permits real-time video-rate image reconstruction on a recent computer.

This equation, combined with Equ. 4 and the description of the grid pattern as a discrete series of on and off lines of chosen widths, permits direct numerical calculation of the image intensity as a function of defocus, and to extract the intensity at focus and the depth of focus (Fig. 4). As for LC, however, it is interesting to derive a simpler analytical expression that permits relating the system parameters to its performance. We will assume that the projected pattern is a 1D sine pattern with period *p*, a reasonable assumption for small values of *p* for which apodization by the optical system efficiently cuts out harmonic frequencies. Using, again, the approximated shape for the illumination beam of Equ. 6 et 7, it is straightforward to obtain that the transmission of such pattern through the optical system is exp 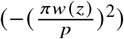. As a result, the same transmission factor applies to the SIM image whose intensity can be written as:

**Fig. 4.**
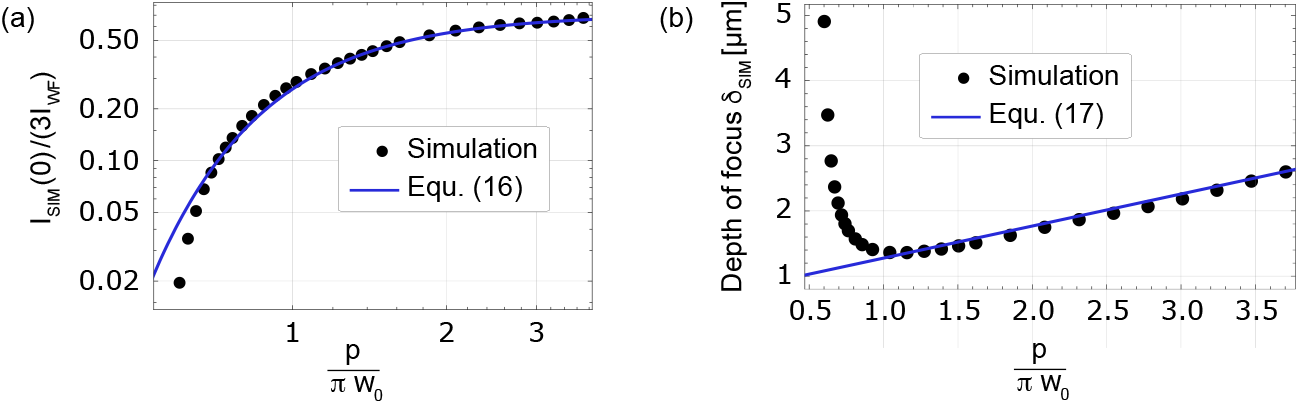
Structured illumination sectioning in coherent reflection microscopy: numerical and analytical analysis. (a), calculated normalised intensity as a function of *p* normalized to *πw*_0_ (black dots), and theoretical predictions from Equ. 16 (blue line) showing excellent agreement. Both are normalised to the full field case with equivalent illumination, corresponding to I_1_ I_2_ I_3_. (c), simulated depth of focus as a function of *p* normalized to *πw*_0_ (black dots), and theoretical predictions from Equ. 17 (blue line) in agreement for 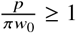. Analytical predictions for smaller values diverge from the exact calculation that reaches a minimum for *p* = *πw*_0_.

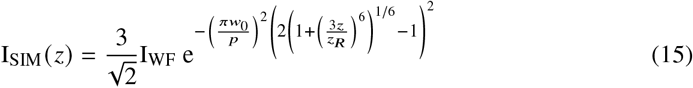

Here I_WF_ is the intensity of the widefield image obtained by averaging the three patterned images I_1_, I_2_ and I_3_. From equation 15, we obtain the SIM image intensity at focus I_SIM_ (0) and the axial resolution *δ*_SIM_:

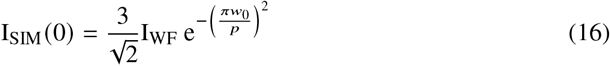

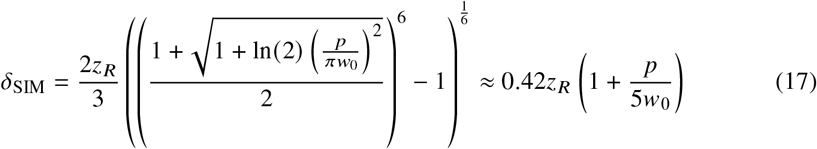

As for the LC case, these theoretical formula permits linking straightforwardly the system parameters to its performance. However, comparing with the more accurate numerical calculation, we observe that while the SIM image intensity is well described by equation 16, the linear variation of *δ*_SIM_ with the period *p* is only appropriate for large enough periods (Fig. 4): indeed, it is well known that optimal sectioning is achieved at a grid frequency corresponding to half of the support of the transfer function of the system, a behaviour that is not recapitulated by our analytical approach. This is easily understood when considering that this optimal grid frequency derives from the presence of side lobes in the real point spread function, which we have neglected in the Gaussian-Lorentzian approximation. The range of validity of equ. 17 is therefore restricted to *p* > *πw*_0_. Note, however, that this corresponds to the useful part of the curve that results in a non negligible image intensity while maintaining optical sectioning close to its optimal value.

With these precautions, we can deduce an optimal spatial period for the illumination pattern that provides efficient optical sectioning while avoiding excessive reduction in image intensity. In the case of SIM, we obtain *p* ≈ *πw*_0_ ≈ λ/INA, a value for which I_SIM_(0) ≈ 0.26I_WF_ and *δ*_SIM_ ≈ 0.68𝓏_*R*_. With our simulation parameters, this corresponds to *δ*_SIM_ ≈ 1.6 *µm*.

### 3.3 Comparison with experimental measurements

As a next step, we have validated the analytical descriptions above by comparing their predictions with experimental values: to this aim, we have used a coverslip coated with a thin (≈5 nm) layer of gold and covered with objective immersion oil, providing a reflective interface between two media of similar refractive indices (Fig.5(a)). For both LC and SIM, we have systematically measured the reflected intensity profile as a function of axial position, for different numerical apertures, magnifications and refraction indices.

**Fig. 5.**
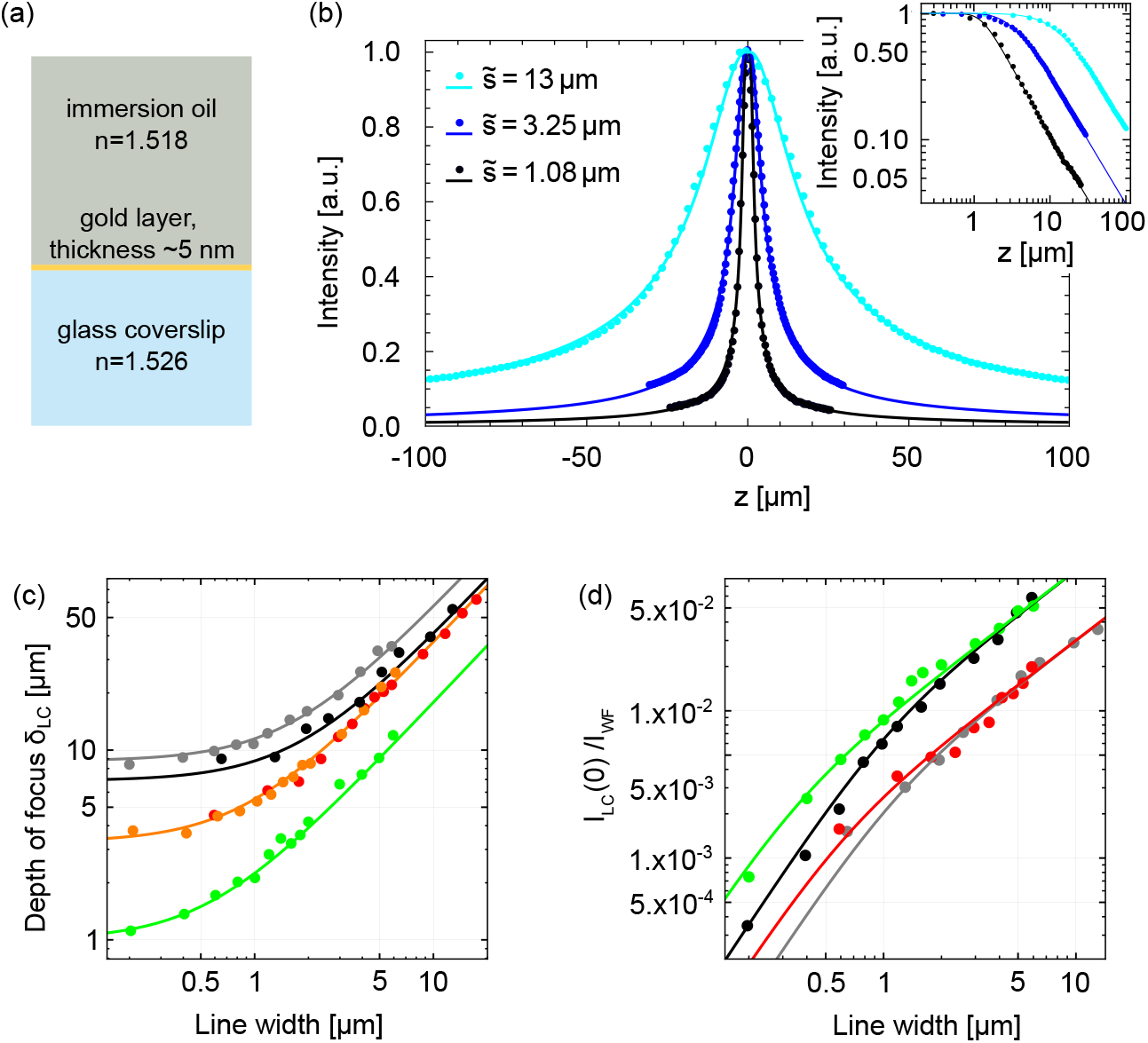
Line confocal detection in coherent reflection microscopy: experimental results compared with analytical predictions. (a), scheme of the test sample used for measuring the reflection intensity profile. The gold layer acts as a thin, homogeneous reflective interface between two media of similar refractive indices. (b), experimental intensity profile reflected as a function of the distance from the focal plane (dots), and corresponding theoretical curve numerically estimated from Equ. 11 (solid lines). Curves were obtained for INA=0.46 and varying 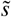 values. Inset, same curves plotted in log-log scale emphasizing the 𝓏^−1^ dependence of the LC image intensity away from the focus. (c), experimentally measured FWHM axial sectioning values as a function of line width 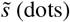, compared with analytical predictions of Equ. 13 (solid lines). Curves were obtained with a 60x, 1.35 NA, oil immersion objective (INA=0.46, orange; INA=0.93, green, INA=0.33, gray), a 20x, 0.85 NA, oil immersion objective (INA=0.46, red) and a 20x, 0.75 NA, air immersion objective (INA=0.3, black). (d), reflected intensity at focus from experimental data (dots) and analytical predictions (solid lines), for the same conditions as in (c). A constant readout noise from the camera was measured independently and subtracted from the raw data.

First we have focussed on the line confocal case: experimental data confirm the square-root-of-Lorentzian intensity dependence with the axial position (Fig. 5(b)), both close and further away from the focal plane. We subsequently quantify the FWHM width of this profile. Results obtained for three objectives with different magnifications, numerical apertures and immersion media (oil or air) are in excellent quantitative agreement with the theoretical prediction for *δ*_LC_ (Fig. 5(c)). We also reproduce quantitatively the prediction for the sectioned image intensity normalized to that of the wide field image (Fig. 5(d)).

Of note, even a INA of 0.93, largely exceeding the limit of the paraxial approximation, still yields results in quantitative agreement with our simple analytical formulas (Equ. 11 and 13). This illustrates the benefit of these simple formulas for the description of the performance of line confocal systems. They can, for example, help establishing the optimal parameters of the imaging system that can be tuned depending on the sample or process under study.

Similarly, we have measured experimentally the optical sectioning obtained with SIM. Figure 6(a) shows a comparison between experimental I_SIM_ (𝓏) profiles (green dots); numerical calculations obtained from the combination of Equ. 4, Equ. 14 and the binary description of the grid pattern as a series of alternating bright and dark illumination lines (solid gray line); and theoretical curves from Equ. 16 (black dotted line). The curves were obtained using or simulating a 60x, oil objective with INA=0.46 and *p* = 1.3 *µ*m (corresponding to 3 on lines alternating with 3 off lines on the DMD): both the numerically calculated and the analytical curves describe the data well, although the simplified theory does not include the secondary maxima around the focus. As outlined in the previous subsection, this results from the approximation of the PSF by a Gaussian profile which also induces an incorrect prediction of the sectioning power of SIM for small *p* values.

**Fig. 6.**
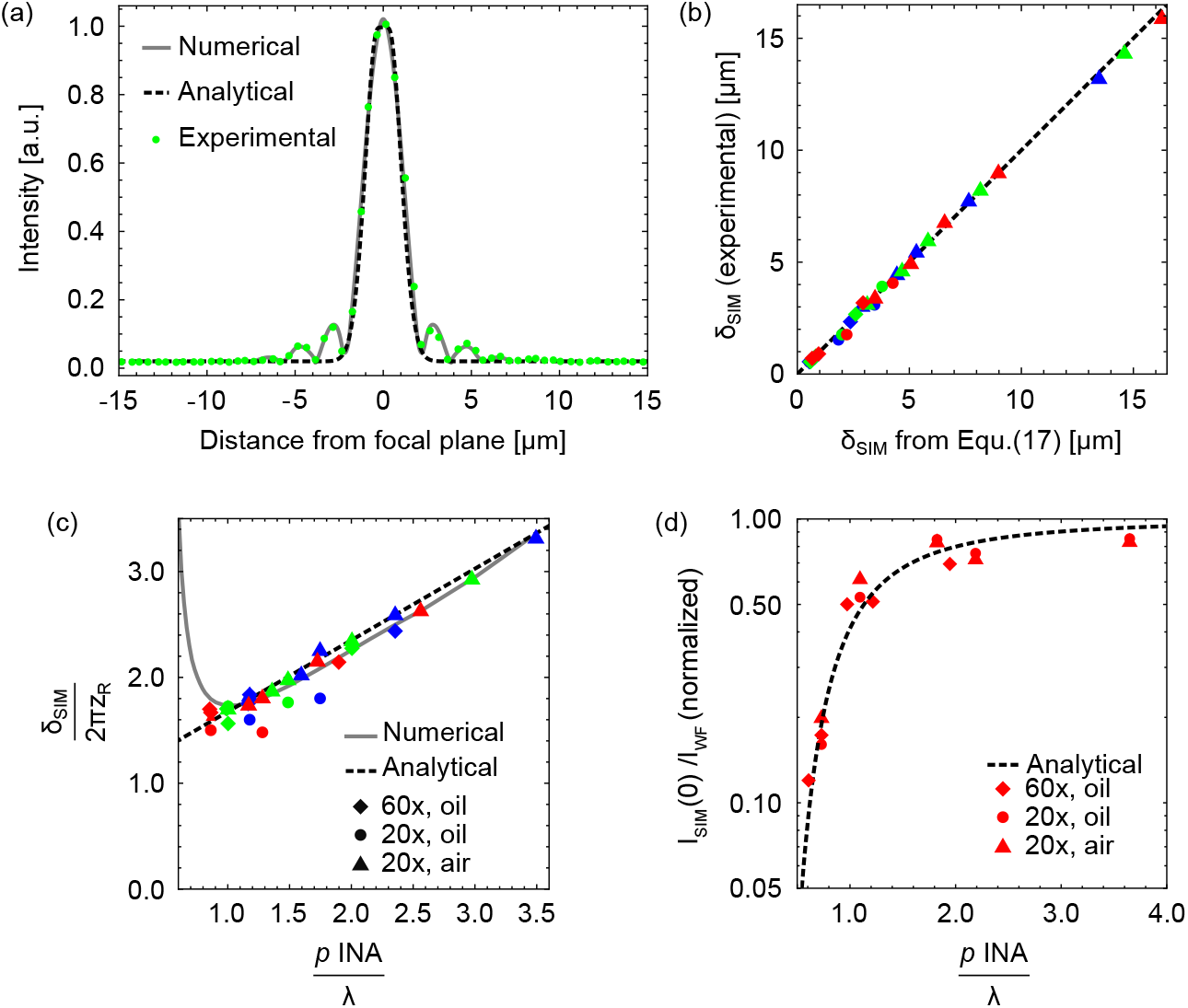
Structured illumination sectioning in coherent reflection microscopy: experimental results compared with analytical predictions. (a), experimental intensity profile from the same sample as in Fig. 5 as a function of the distance from the focal plane (dots), obtained with INA = 0.46 and *p* = 1.3 *µ*m ; corresponding theoretical curve numerically estimated from Equ. 4 and Equ. 14 (solid line, grey) or calculated from Equ. 15 (dotted line, black). (b) and (c), experimental depth of focus (solid symbols) for various objectives (different symbol shapes, legend in panel (c)), INAs (ranging from 0.21 to 0.93), values of *p* (ranging from 0.65 to 3.9 *µ*m) and wavelengths (blue, 453 nm; green, 532 nm; red, 635 nm). In (b), experimental values are plotted against analytical predictions. The black dotted line corresponds to *y* = *x*. In (c), the same values normalized to 2*π*𝓏_*R*_ are plotted against 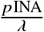, and compared to analytical and theoretical predictions. (d), reflected intensity at focus from experimental data (dots) and analytical prediction (black dotted line), for the same conditions as in (b,c) and a wavelength of 635nm.

This simplified approach nevertheless proves relevant for practical cases, where larger *p* values are generally used to maximize the contrast of the grid and the brightness of the sectioned image. As illustrated in Fig. 6(b), Equ. 17 describes accurately the depth of focus achieved experimentally for the practical cases that we have implemented, from 0.5 to over 15 *µ*m. Indeed, all these cases correspond to 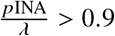, and do not cover the region where numerical and theoretical predictions diverge strongly that presents little practical interest. This is illustrated on Fig. 6(c) that presents the same data as in Fig. 6(b), but normalized such that they can be compared to the two predictions. Comparing Fig. 6(b) and (c) outlines that the greatest part of the amplitude in *δ*_SIM_ variation is linked to the INA (through 𝓏_*R*_) rather than to *p* over a fairly wide range. This is in contrast with the sectioned image intensity, shown on Fig. 6(d): here, the benefit of using patterns with sufficiently large periods appears clearly, in particular around 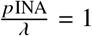. Fig. 6(d) also demonstrates the good agreement between experimental data and Equ. 16.

Together, experimental measurements confirm the validity of our analytical approach that provides simple relations between the setup parameters - wavelength, INA, illumination line width or period - and the performance of the sectioning both in terms of depth of focus and of image brightness.

### 3.4 Noise and spurious signal in optically-sectioned coherent reflection microscopy

In this subsection, we now focus on the quality of reflection images achieved with or without these optical sectioning approaches. Indeed, the (random) shot noise and remaining spurious signal in the final image determine respectively the precision and accuracy of the quantitative information that can be retrieved from images and are critical parameters to determine which method should be used to suppress the incoherent signal from coherent reflection images. Here, we have considered three possible approaches for the removal of the incoherent background: in addition to LC and SIM, subtraction of a known background to the WF image is also considered as a means to recover an unbiased image. Finally, as quantified in Fig. 3(b) and 5(d), LC images are much dimer than their WF counterpart: we have therefore considered two practical cases for LC, one in which the illumination intensity equals that of the WF case, and one in which the illumination intensity can be increased to exploit the full bit depth of the camera (assuming that the WF image and the modulated images for SIM also use this full bit depth).

We have considered different sources of incoherent background corresponding to different practical cases. First, to assess the precision of the final image, two situations are modelled: (i) the case of a homogeneous reflective plane located far from the focal plane; and (ii) the case of diffuse reflections evenly distributed above the focal plane. Secondly, we consider accuracy achieved in the presence of (iii) a homogeneous background and (iv) a structured background, both stemming from structures located in a single plane in the vicinity of the focal plane.

Using Equ. 11, 13, 16 and 17, we compute the signal-to-noise ratio (SNR) of the final image (see Appendix) for cases (i) and (ii), and the remaining error for cases (iii) and (iv). The former evaluates of much random noise is present in the final image, and hence how precise this image is (e.g., how weak signal variations can be separated from noise), while the latter assesses of much residual incoherent signal is present, and hence how accurate the final image is (e.g., is the average intensity biased, or do artifacts from outside the focal volume appear on images?).

The results are presented in Fig. 7: we first compare the experimental intensity profiles from a single reflective plane for LC and SIM (Fig. 7(a)), that illustrate the much slower decay of the LC signal with the distance from this plane: as a result, structures located in the vicinity of the focal plane will be suppressed from the LC image with a much lesser efficiency than from their SIM counterpart. On the other hand, the SIM profile saturates rapidly to a final value corresponding to the standard deviation of the shot noise on the three modulated images used to compute their sectioned counterpart, such that a significant residual random noise is always present on SIM images even in the case of spurious signals stemming from structures located far from the focal plane.

**Fig. 7.**
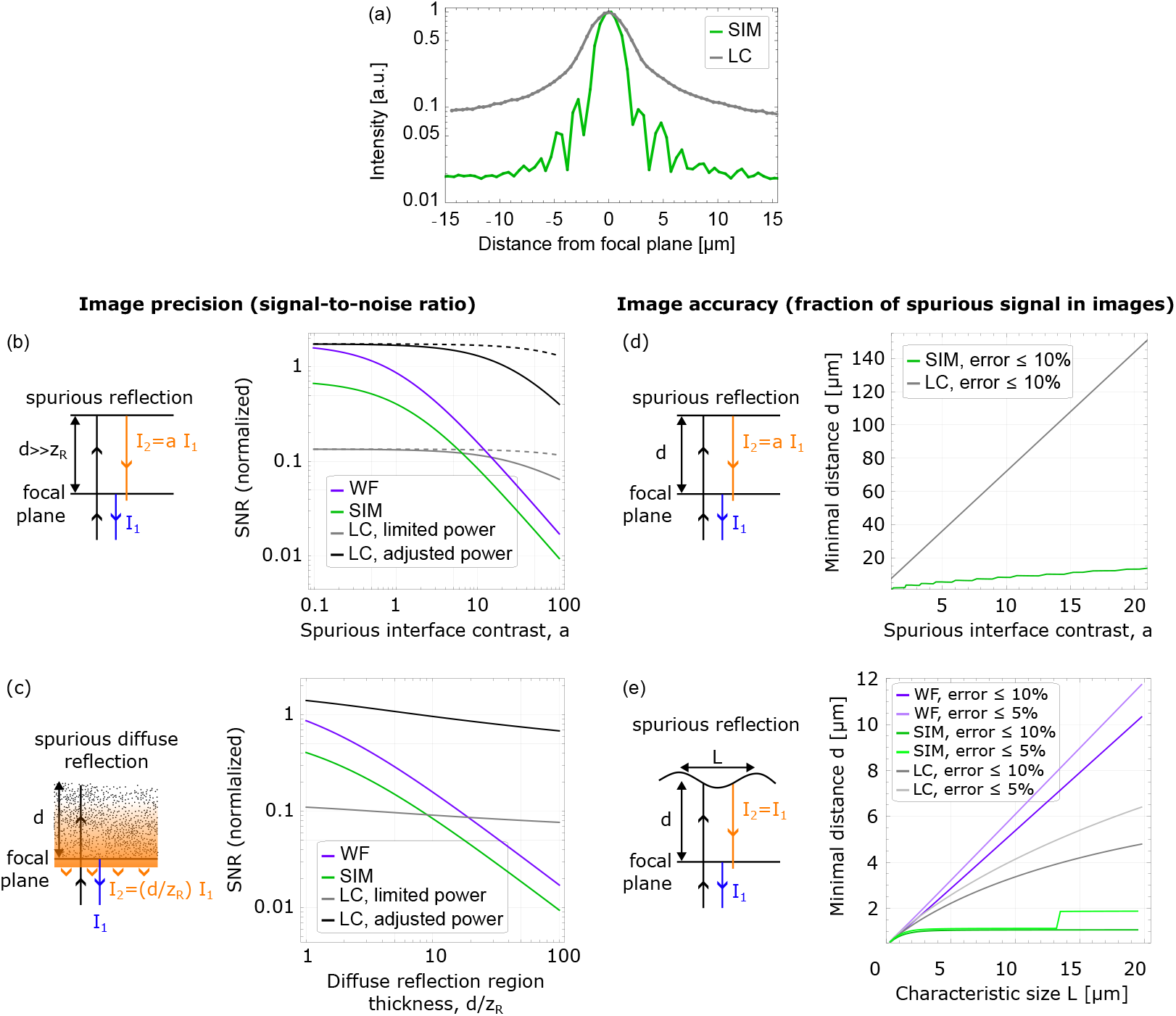
Accuracy and precision of coherent reflection images achieved by optical sectioning. (a), typical experimental intensity profile from a homogenous reflective plane as a function of the distance from the focal plane for LC (gray) and SIM (green). Profiles were acquired using INA=0.46, 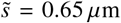 (for LC) and *p* = 1.3 *μ*m (for SIM). (b,c), signal-to-noise ratio (SNR) for the different techniques as a function of the spurious reflection intensity of an interface located far from the focal plane (b) and of the thickness of a diffusing medium (c). In (b), solid lines correspond to *d*/ *𝓏* _*R*_ = 10 while dashed lines correspond to *d*/ *𝓏* _*R*_ = 100. (d,e), minimal distance between the focal plane and interfaces inducing a spurious contrast in the final image, required to reduce the resulting error below a set threshold, for a homogeneous reflective plane (d) and a structured interface with characteristic size L (e). Jumps in the SIM curve stem from the side lobes of the sectioning curve.

These particularities, in turn, translate into benefits or disadvantages for suppressing different types of spurious signals from reflection images. Fig. 7(b) and (c) quantify the SNR in the presence of a strong, distant homogeneous reflection or a thick diffuse background, respectively (see Appendix for the detailed calculations and normalisation): in both cases, WF (with background subtraction, purple lines) and SIM (green lines) performance drop sharply with the increasing amount of spurious intensity, because both methods rely on the a posteriori removal of the spurious signal that is still acquired on the camera, hence progressively reducing the effective bit depth allocated to the in-focus signal when using the full bit depth of the camera. This effect is well known in optical coherence tomography (OCT), a technique for which it is the main limitation for in-depth imaging inside biological tissues. Interestingly, SIM performs even worse than WF with background subtraction, because the noise from the three combined images adds up in the resulting image. Note, however, that background subtraction requires accurate knowledge of the background intensity, which might be difficult to achieve.

SNR in the LC image, on the other hand, depends only weakly on the intensity of very distant spurious reflections because they are mostly cut out by the detection process. Because the acquisition time per pixel is greatly reduced, however, SNR at the same illumination power is lower than WF and SIM as long as the spurious intensity is not much greater than that from the focal plane (gray lines). On the contrary, when considering the case where the illumination power can be increased to exploit the full bit depth of the camera, we find that the SNR of the LC image is then constantly larger than that of other methods (black lines).

This analysis indicates that provided that enough illumination power is available to compensate for the shorter exposure time per pixel, LC is a method of choice to remove the background stemming from a distant interface, and that if the background can be accurately estimated, WF imaging should be preferred to SIM and even to LC if excitation power is limited. For intermediate, unknown background intensities, SIM can still be beneficial if the illumination power is limited.

The picture is different when considering the accuracy of the final image in the presence of reflecting structures in the vicinity of the focal plane (Fig. 7(d) and (e)): here, we evaluate how much unwanted incoherent signal is left in the final images and can bias the quantitative interpretation of the interference pattern. Setting a maximum acceptable fraction of spurious signal, we calculate the minimal distance between the spurious reflective structure and the focal plane that permits sufficient sectioning to achieve this set accuracy, as a function of the spurious structure contrast (Fig. 7(d)) or of its characteristic size (Fig. 7(e)): in both cases, SIM performs significantly better than LC due to the faster decay of its sectioning curve (Fig. 7(a)). While WF can, in theory, suppress a constant background irrespective of its distance from the focal plane provided that its amplitude is known, the approach fails as soon as unknown structures appear in the background.

Contrary to the case of a distant interface, we thus find that the best approach to minimize artifacts in images acquired on complex samples, for which significant modulation can stem from out-of-focus structures located only a few microns away from the focal plane, is SIM rather than LC, outlining the complementarity of the two sectioning approaches.

We illustrate this conclusion experimentally in Fig. 8 on two typical application examples of RIC microscopy: thin organic film mapping, and cell adhesion characterization. Fig. 8(a) presents images of a 6-nm-thick patterned brush of poly-(N-isopropylacrylamide) (PNIPAM). Dense, thin PNIPAM brushes act as barrier for cell adhesion by preventing protein adsorption, and fibronectin- or collagen-covered micrometric patterns devoid of brush can be used to force cell adhesion onto reproducible shapes [30] or control mobility on surfaces [18]. To inspect such sample during preparation, air objectives can be beneficial at intermediate steps as they do not contaminate the sample with immersion oil, but a strong reflection then arises from the air/glass interface at the bottom of the functionalized coverslip, typically 10 times more contrasted than the interface under study.

**Fig. 8.**
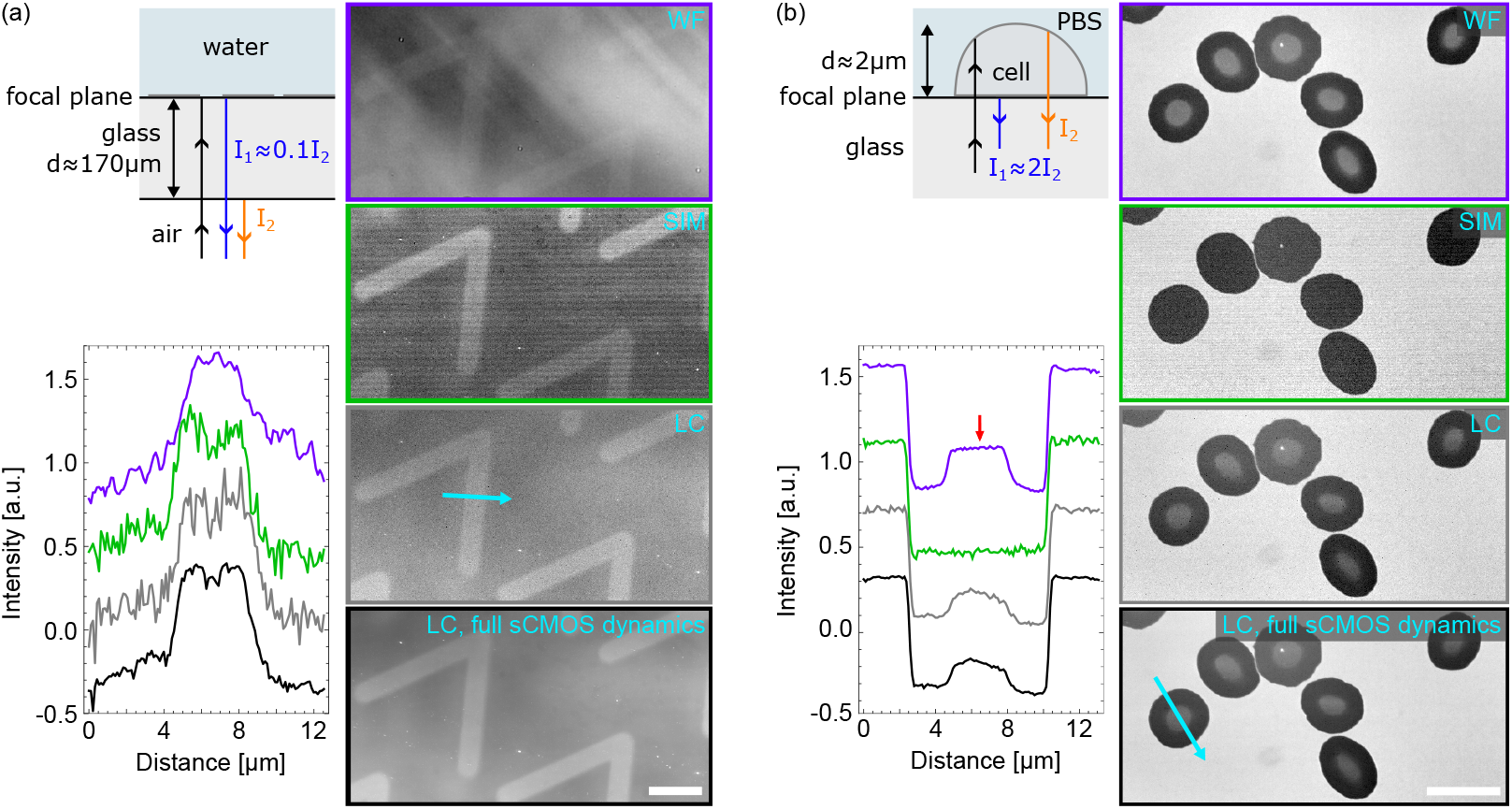
Accuracy and precision of optically sectioned coherent reflection images: experimental RIC examples. (a), imaging of a patterned PNIPAM brush using WF, SIM and LC RIC microscopy. All images are adjusted for best visibility of the brush pattern, effectively removing a constant offset and normalizing the image intensity. Intensity profiles along the blue arrow are shown on the left. Profiles were all normalized to allow direct comparison of the noise, with an arbitrary offset to improve readability. Images were obtained with a 20x, 0.75NA air objective with INA=0.46, *λ* = 532 nm, 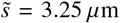 (for LC) and *p* = 3.9 *μ*m (for SIM). (b), Red blood cells adhered on a fibronectin-coated coverslip imaged with WF, SIM and LC RIC microscopy. Images and intensity profiles are offset and normalized as in (a). The red arrow points to the spurious signal reflected from the cell upper surface. Images were obtained with a 60x, 1.35NA oil objective with INA = 0.58, *λ* = 532 nm, 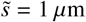 (for LC) and *p* = 1.3 *μ*m (for SIM). All scale bars are 10 *μ*m.

Testing the same four options as presented above to remove this incoherent background, we find that, using the full bit depth of the camera, although subtraction of an offset in WF imaging permits to distinguish the micropatterns, imperfect knowledge of the background results in residual spurious intensity variations of amplitude larger than the contrast of the patterns. In contrast, SIM permits to distinguish clearly the latter, although noise on the image and a small residual modulation from the fringe pattern are visible. LC with the same excitation intensity provides a result similar to SIM (as expected from the calculations in Fig. 7(b)), but clearly outperforms other options when using the full bit depth of the camera. Because excitation power is limited in our setup, this was achieved experimentally by summing several successive images such that electronic noise from the camera is slightly stronger than it should.

We then considered the case of adhered red blood cells on a fibronectin-covered coverslip. Such immobilization method is useful to probe the rheology of the cells, e.g. using atomic force microscopy [19], but interpreting the results requires knowledge of the adhesion area which controls membrane tension and the cell geometry. Here, at high fibronectin density, cells lose their biconcave shape and flatten on the coverslip to maximize adhesion energy, adopting a lens shape due to increased surface tension. As displayed on Fig. 8(b), the tight contact area appears dark on RIC images due to the higher refractive index inside cells compared to that of the surrounding medium (phosphate buffer saline, *n* ≈ *n*_water_). WF RIC images, however, show a brighter elliptical area in the center of the adhesion patch: this artifact stems from the reflection of light on the ellipsoidal upper surface of the cells that refocus the light onto the focal plane, producing an incoherent image of the aperture diaphragm superimposed with the interference image. Such artifact can bias the interpretation of the adhesion pattern if it is mistakenly assimilated to an increasing distance between the cell membrane and the glass surface.

Using SIM solves this ambiguity by efficiently removing from the RIC image the out-of-focus halo created by the cell upper surface. While this comes at the cost of a slightly enhanced random noise in images, quantitative interpretation of images and subsequent processing (e.g., to compute the adhesion area per cell) is greatly facilitated. LC, on the other hand, does not completely suppress this artifact because it arises from a structure very close (2-3 *μ*m) to the focal plane: while the halo is slightly dimmed, the decrease in transmission with the distance from the focal plane is too slow to prevent transmitting a significant part of it (see e.g. Fig. 7(a)). LC images, in particular when using the full bit depth of the camera, appear smoother and exhibit reduced random noise, but this increased precision comes at the cost of a reduced accuracy that, contrary to precision, cannot be improved by averaging of successive images.

## 4. Discussion and conclusion

In this article, we have derived equations to describe the performance (image intensity, depth of focus) of LC and SIM for optical sectioning in coherent reflection imaging. This analytical approach, while relying on simplifying assumptions, is validated in a variety of practical cases through systematic comparison with experimental data. It is well known that interpreting RIC signals requires to carefully take into account corrections due to changes in reflection coefficients, polarisation or refraction effects at high INAs [15, 31]. We have shown here, in contrast, that a simplified approach yields quantitatively correct predictions of the performances of the sectioning in coherent reflection imaging (including RIC imaging). As a result, equations 11, 13, 16 and 17 can be used to simply and reliably design the optical sectioning part of the optical setup depending on the type of out-of-focus incoherent background that is expected, and in particular whether it stems mostly from structures close to (for which SIM performs best) or distant from (for which LC is preferable) the focal plane.

As outlined at the beginning of this paper, while developed using assumptions derived from the field of reflection interference microscopy, our analytical approach for performance evaluation also applies for SIM or LC sectioning in other microscopy techniques such as widefield reflection microscopy (e.g. in fields such as ophtalmography, where bright out-of-focus reflections far from the focal plane from the cornea limit the contrast of the structures of interest [32, 33]), optical coherence tomography (OCT) [34] or, with appropriate changes due to incoherent imaging, to fluorescence imaging. Indeed similar observations concerning the relative benefits of LC and SIM have already been partly discussed: implemented using sufficient illumination power, LC permits allocating the whole camera bit depth to imaging of the structures of interest [35] while SIM performs more poorly on thick and scattering tissues [36].

Other camera-based approaches to fast full field imaging with optical sectioning exist in the literature that were not considered in the present study, notably relying on the parallelization of confocal microscopy. Multipoint-scanning confocal [37] and spinning disk confocal [36] both use an array of point sources created by pinholes that are scanned across the image to increase the frame rate compared to confocal microscopy by a factor equal to the number of illuminating points. While such systems are very efficient in a number of applications, their complexity and lack of flexibility make them challenging to adapt to reflection imaging. For example, commercial spinning disk modules cannot incorporate easily polarization optics or INA control features. We therefore could not compare their experimental performance with that of LC and SIM, and a detailed analysis of these is not present in the literature. Briefly, however, one can convince oneself that they represent intermediate compromises between SIM and LC: their rejection of signal stemming from structures close to the focal plane, identical to that of confocal, is excellent and compares to SIM. However, similar to SIM, out-of-focus background is more poorly rejected than in confocal microscopy or LC, with a transmitted fraction of out-of-focus background equal to the surface coverage of the scanning points [37, 38]. This is because contrary to LC, the camera here acquires the signal continuously on all pixels, instead of synchronizing illumination and acquisition on each pixel. Improving the rejection requires spreading out the scanning point, thereby decreasing the signal intensity and frame rate, up to the single point case corresponding to the classical confocal microscope. As for SIM and unless coupled with other techniques, these approaches thus perform better on relatively thin or weakly reflecting/scattering samples.

In conclusion, the results presented here help rationalizing the choice of an optical sectioning method and provide guidelines to their best use: in the case of a large background stemming from an extended region, or a single structure located far from the focal plane, LC outperforms SIM or background subtraction. On the contrary, when spurious signals arise from structures close to the focal plane, SIM is the best option to efficiently suppress artefacts from images and improve contrast. While (point scanning) confocal or spinning disk coherent reflection imaging could be considered, their complexity and cost is much larger than that of SIM implementation, which is easily accommodated in an existing microscope. Furthermore, both LC and SIM can be implemented using the same optical setup adaptation on a widefield microscope: it can thus be beneficial to develop both, so that the most appropriate technique can be used depending on the sample under study. Note that such solution also permits adding to the setup illumination patterning abilities that can be used for photopatterning of surfaces, or optogenetic.

We hope that this quantitative study of the performance of LC and SIM for coherent reflection microscopy, and in particular the precise comparison of precision and accuracy achieved with different approaches, will help with their pratical implementation on a variety of widefield microscopes.

## Acknowledgments

The authors thank Lionel Bureau for kindly providing the patterned PNIPAM brush sample, Claude Verdier for help with the RBC sample preparation, and Jerome Mertz for fruitful discussions.

## Author contributions

C.V. and D.D. conceived the study, derived the theory and drafted the original manuscript. D.D. and A.N. designed experiments and collected data. D.D. analyzed data, performed numerical analyses, supervised the project, and raised funds. All authors contributed to manuscript review and editing.

## Data Availability Statement

Data underlying the results presented in this paper are not publicly available at this time but may be obtained from the authors upon reasonable request.

## Appendix: derivation of signal-to-noise ratio formula for WF, LC and SIM

In order to compare the shot noise present in the images obtained through different modalities, we need to specify how the illumination intensities compare for these different methods. For the calculation presented here, we have considered the case where the images were acquired using the full bit depth of the camera, such that the average illumination intensity per pixel and per image might differ between WF, SIM and LC. In addition, for LC imaging, we have also considered the common case where the illumination power on the DMD was limited to that of the WF case - a common situation for moderately powerful sources. In this latter case, only a fraction of the full bit depth of the camera is used.

Let us call *N* the maximum count per pixel when the full bit depth of the camera is used. We assume that the signal variation on each pixel arises from the shot noise, and is thus proportional to the square root of its average value: *N* is thus such that 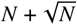 is equal to the full bit depth (e.g., 256 for a 8-bit camera). We will neglect the electronic/readout noise as it is normally negligible compared to *N* for scientific cameras. If a given image results from the incoherent addition of an in-focus signal of intensity *S* and an out-of-focus signal of intensity *B*, we thus get *S* + *B* = *N* and the total noise on the image is 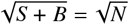. In addition, since 3 widefield images are required to obtain a SIM image, we will compare the noise in the resulting SIM image with that of 3 averaged WF or LC images to keep the acquisition time constant and present a fair comparison.

### Case 1: spurious signal from a single reflection far from the focal plane

We first consider the case depicted in Fig. 7(b): a single reflection located at a distance *d* >> 𝓏 _*R*_ from the focal plane, with a reflection coefficient *a* times that of the focal plane structure. The WF image has an intensity *N* = *S* + *B* = *S*(1 + *a*), and thus a noise 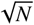. Averaging three WF image thus yields the signal-to-noise ratio:

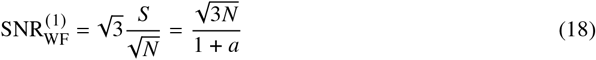

In the case of LC, we consider the case of optimized parameters for which 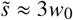, corresponding to *S* ≈ 0.006 *S*_WF_. In the limit *d* >> 𝓏 _*R*_, combining equations 7 and 10 yields 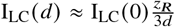 so that 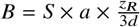. If the illumination intensity is not adjusted, we get for the total image intensity:

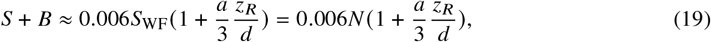

where *S*_WF_ was taken here at its maximum value as the low LC intensity doesn’t require adjustment to avoid saturation. Thus:

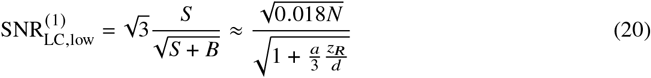

If, on the contrary, the illumination intensity is adjusted to use the full bit depth of the camera, we obtain 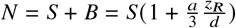 and:

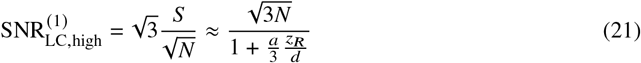

In the case of SIM, we should consider the intensity and noise on each of the three modulated images I_1_, I_2_ and I_3_. Each of these images has an average intensity *S* + *B* and a maximum intensity *S*(1 + *m*) + *B*, with *m* the modulation depth induced by the grid pattern. We thus have *B* = *a* × *S* and *N* = *B* + *S*(1 + *m*), hence *S* = *N*/(1 + *m* + *a*). The average SIM image intensity in the absence of noise is thus 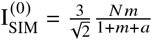. To calculate the noise on the reconstructed image, let us write each of the modulated images I_1_, I_2_ and I_3_ as:

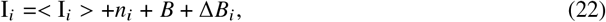

with *i*=1,2 or 3. B is the average intensity from the spurious reflection and *n*_*i*_ and Δ*B*_*i*_ the noise in image *i* on the signal originating from the focal plane and from the spurious reflection, respectively. Due to the stochastic nature of shot noise, these quantities can be described by the following equations:

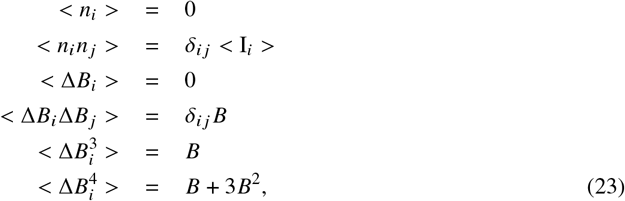

where *δ*_*ij*_ is the Kronecker delta function. Because of the nonlinear nature of the SIM reconstruction algorithm, obtaining an analytical formula for the signal-to-noise ratio in images is more challenging than in the case of WF and LC imaging. We therefore introduce the approximation that the noise induced by *n*_*i*_ and Δ*B*_*i*_ is small in comparison with the image average intensity, an approximation that fails at high noise level but actually proves surprisingly accurate up to low SNRs. Using this approximation, we now write the squared SIM intensity as:

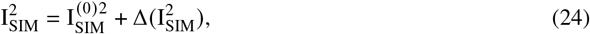

with 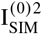 the squared SIM intensity in the absence of noise, and 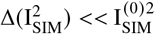. The variance of the SIM intensity due to noise thus writes:

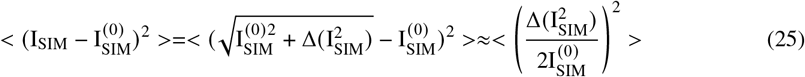

Using equations 23 and 25, we obtain through lengthy but straightforward manipulations:

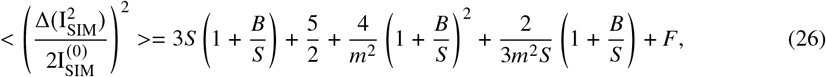

With *F* an oscillatory term that spatially averages out to zero. The assumption 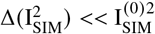 implies that *S* >> *B* >> 1, such that finally:

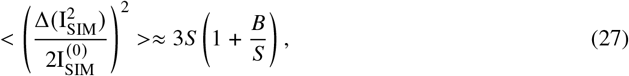

from which we deduce:

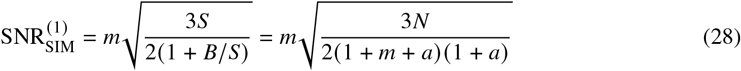

While this equation was obtained in the case of a large SNR, comparison with numerical simulations show good agreement down to SNR ≈ 0.1, thus covering a very large range of experimental situations. In Figures 7(b) and (c) we have considered the nearly-optimal case of *p* = 2*πw*_0_ such that 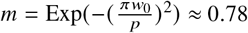. The SNR formula obtained above differ from that of Hagen et al., which we believe is incorrect on the basis of comparison with numerical simulations (not shown) [39]

### Case 2: spurious signal from a collection of small diffusers

We next consider the case of the spurious signal created by a diffusing medium, modelled as a homogeneous distribution of small diffusers each reflecting a small fraction of the incoming light (Fig. 7(c)). The reflection per slice of medium of infinitesimal thickness is assumed to be the same in the focal plane and further away from it, such that the reflected light is a function of the thickness : 𝓏 _*R*_ for the signal, *d* for the spurious reflection. The WF image has an intensity *N* = *S* + *B* = *S* (1 + *d*/𝓏_*R*_), and thus a noise 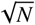. Averaging three WF image thus yields the signal-to-noise ratio:

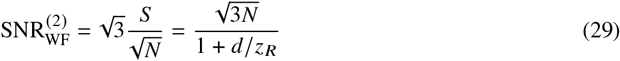

In the case of LC, we again consider the case of optimized parameters for which 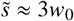, corresponding to *S* ≈ 0.006 *S*_WF_ = 0.006*N*. In the limit *d* >> 𝓏 _*R*_ and using 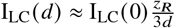, we compute:

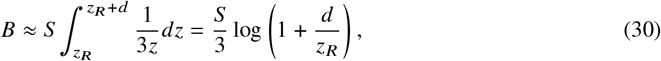

and thus:

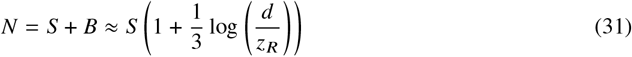

If the illumination intensity is not adjusted, we finally get for the SNR:

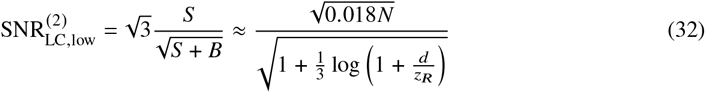

If, on the contrary, the illumination intensity is adjusted to use the full bit depth of the camera, we obtain:

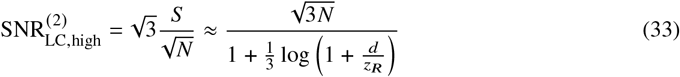

Finally, in the case of SIM, as for case 1 we have *N* = *S*(1 + *m*) + *B* = *S*(1 + *m* + *d*/𝓏 _*R*_), and thus:

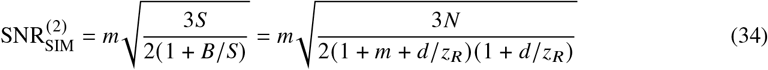

